# Oxytocin receptor Gαi signaling drives valence reassignment to facilitate reversal of social trauma-induced avoidance

**DOI:** 10.64898/2026.08.08.743644

**Authors:** Theresa Schäfer, David Keller, Francisco de los Santos, Anna Bludau, Anna Krinner, Tatiana Korotkova, Inga D. Neumann, Rohit Menon

## Abstract

Traumatic social experiences often generate social deficits that form the core symptom of psychopathologies like social anxiety disorder (SAD). The failure to overwrite these aversive social associations is a fundamental barrier to recovery from SAD. Using the mouse social fear conditioning (SFC) paradigm, we identify a discrete population of oxytocin receptor (OXTR)-expressing neurons in the caudal lateral septum (LSc^OXTR^) that act as a critical hub for adaptive learning during social fear extinction. Using calcium imaging in mice undergoing extinction training, we show that successful valence reassignment is best predicted by the precise temporal inhibition of LSc^OXTR^ neurons during the contact phase of social investigation. This contact-associated suppression of LSc^OXTR^ neurons was absent in mice that failed to extinguish fear. Notably, successful extinction was also associated with enhanced oxitocinerg innervation within the LSc. Chemogenetic silencing of LSc^OXTR^ neurons prior to extinction impaired extinction, indicating that a specific temporal activity pattern of LSc^OXTR^ neurons is necessary for adaptive learning. Pharmacological dissection of downstream OXTR signaling revealed that selective activation of Gαi-coupled, but not Gαq-coupled, signaling accelerated extinction and promoted social approach. Our findings establish that extinction success is governed by the temporally precise, Gαi-mediated silencing of LSc^OXTR^ neurons during social contact, revealing a novel mechanism for adaptive social valence processing with direct implications for treatment-resistant SAD.

## Introduction

The precise orchestration of social behavior is fundamental to our well-being and survival. Hence, traumatic social experiences that disrupt adaptive social functioning can be extremely debilitating. Social fear and avoidance, which are often caused by social trauma, are core symptoms of psychopathologies like social anxiety disorder (SAD) ^1^. The therapeutic state-of-the-art for SAD consists of cognitive behavioral therapy (CBT), including exposure therapy, that drives the reassociation of a positive emotional valence with social cues ^2^. However, remission rates for CBT are approximately 50% ^3, 4^, indicating that a significant subpopulation of SAD patients is treatment-resistant. Animal models, such as the social fear conditioning (SFC) paradigm, have been shown to elicit robust social avoidance in both male and female mice ^5, 6^. The extinction phase of the SFC paradigm is akin to exposure therapy and generates two groups of mice: (i) those that successfully respond to extinction training and, as a result, do not express social fear anymore (Responders; Res), and (ii) those that do not respond to extinction training and still display social fear thereafter (Non-responders; NRes) ^7^. Thus, leveraging such preclinical models to dissect the neuronal basis of individual differences in CBT success is essential to identifying novel avenues for SAD treatment.

The lateral septum (LS) is a subcortical brain structure that regulates a wide range of social behaviors ^8, 9^, including the extinction of social fear ^6, 10–13^. Local neuromodulation by the neuropeptide oxytocin (OXT) is critical in this regard ^6, 10^. OXT reaches the LS via long-range projections from the supraoptic (SON) and paraventricular nuclei, and its enhanced release within the LS has previously been linked to reduced social fear expression during extinction ^6, 13, 14^. This social fear-reducing effect of OXT is mediated via OXTR-expressing LS (LS^OXTR^) neurons. In vitro studies have shown that binding of OXT to OXTR primarily activates signaling via the Gαq pathway, while OXT can also drive the coupling of OXTR with Gαi proteins to transduce different, context- and cell-type-specific intracellular signals ^15, 16^. In neurons, activation of Gαq signaling enhances excitatory postsynaptic potentials, whereas Gαi signaling suppresses neuronal activity ^17, 18^. The conventional understanding of how canonical OXTR signaling influences behavior centers on its coupling with Gαq proteins ^15, 19^. However, recent evidence for OXTR-Gαi signaling modulating fear responses challenges this hypothesis ^20^. In the context of social behavior, the precise neuromodulatory mechanism by which LS^OXTR^ neurons drive valence association and regulate adaptive responses to aversive social stimuli remains unknown.

In this study, we dissect the temporal and mechanistic underpinnings of LS^OXTR^-mediated regulation of valence association that occurs during social fear extinction. Single-cell Ca^2+^ imaging using miniscopes showed that LS^OXTR^ neurons were excited in response to aversive social stimuli and silent in response to appetitive social stimuli. Further temporal dissection of Ca^2+^ dynamics reveals that LSc^OXTR^ neurons are inhibited specifically during the contact phase of social investigation when the stimulus is associated with a positive valence. This contact-associated inhibition, followed by post-contact excitation, distinguishes mice that successfully extinguish social fear (i.e., Res mice) from those that do not (i.e., NRes mice). Chemogenetic inhibition of LSc^OXTR^ neurons disrupts this temporally structured activity pattern and impairs social fear extinction, demonstrating that the transient silencing of these neurons during social contact is necessary for adaptive social learning. Finally, pharmacological manipulation of local OXTR-Gαi signaling facilitated the extinction of social fear, indicating a novel neuromodulatory role for Gαi signaling in LS^OXTR^ neurons in driving adaptive learning during social fear extinction and alleviating the effects of social trauma.

## Materials and Methods

### Animals

All studies were conducted according to the Guide for the Care and Use of Laboratory Animals of the National Institute of Health and ARRIVE guidelines for animal research ^21^ and were approved by the Government of Oberpfalz, Germany. Wildtype male CD1 mice were purchased from Charles River Laboratories (Sulzfeld, Germany). Transgenic mice expressing Cre recombinase in OXTR-expressing neurons (OXTR-Cre; CD1 background; MGI: 5311702; ^22^), were obtained from the Max-Planck-Institute for Psychiatry (Munich; Dr. Jan Deussing). OXTR-Cre mice were crossed with mice expressing floxed td-Tomato reporter protein (Ai9; C57Bl6 background; MGI: 3809523) to produce OXTR-Cre:Ai9 transgenic mice. All transgenic mice were further bred in the animal facilities of the University of Regensburg (Germany). All animals were housed under standard laboratory conditions (12/12 hrs light/dark cycle, lights on at 07:00 a.m., 21 - 23 °C, 55-60 % humidity, food, and water ad libitum) in polycarbonate cages (16 x 22 x 14 cm) until described otherwise. Mouse cages were changed weekly. All experiments were performed between 08:00 and 13:00 and in adult male mice.

### Social exposure

Male mice were exposed to an age-, sex-, and weight-matched conspecific in their home cage for 3 min to assess baseline sociability before exposure to the SFC paradigm. This test was performed prior to Ca^2+^ imaging experiment.

### SFC paradigm

In order to induce the association of a negative emotional valence to a social stimulus, which is expressed in the form of robust social fear, mice were exposed to the SFC paradigm as previously described ^5, 6^. The procedure spanned three days:

#### Day 1 Social Fear Acquisition

Mice were habituated to a conditioning chamber for 30 sec before an empty wire cage (non-social stimulus) was introduced for 3 min. This was replaced with an identical cage holding an unfamiliar, same-sex conspecific (social stimulus). Mice of the non-conditioned group (SFC^-^) were allowed to freely investigate the social stimulus for 3 min. In contrast, social fear-conditioned mice (SFC^+^) received a mild foot shock (mean of 2 shocks/mouse) upon sniffing at/closely investigating the social stimulus. The a*cquisition* session ended after 2 min if no further contact was observed. During Ca^2+^ imaging experiment, social fear acquisition is performed on day 2, as it is preceded by social exposure sessions.

#### Day 2 Social Fear Extinction

Mice were exposed to a series of stimuli in their home cage: three empty cages (non-social stimuli; ns1 – ns3) followed by six different unfamiliar, same-sex conspecifics (social stimuli; s1 – s6). Each exposure lasted 3 min with a 3-min interexposure interval. During Ca^2+^ imaging experiment, social fear acquisition is performed on day 3 and day 4, respectively, as it is preceded by social exposure sessions.

#### Day 3 Social Fear Extinction Recall

Mice were again exposed to the described series of six different unfamiliar social stimuli in their home cage, following the same timing protocol as on Day 2. During Ca^2+^ imaging experiment, social fear acquisition is performed on day 4 and day 5, respectively, as it is preceded by social exposure sessions.

All *social fear* extinction and recall test sessions were video-recorded and scored by an observer blind to treatment groups using Jwatcher ^23^ and BORIS ^24^ software. Non-social and social Investigation times, i.e., times of sniffing and direct physical interaction with the stimulus, were assessed.

### Surgical procedures

Upon arrival, mice were habituated to the new environment for at least 1 week before surgery. Mice received a subcutaneous injection of the analgesic Buprenorphine (0.05 mg/kg, Buprenovet, Bayer, Germany) 30 min before the start of the surgery, were anesthetized with Isoflurane (4% Isoflurane, Abbott GmbH, Germany) throughout the surgical procedure, and mounted in a stereotactic frame (Kopf Instruments, Canada). Eyes were covered with ophthalmic ointment (Bepanthen, Bayer, Germany) to avoid drying. Before cutting the skin on top of the skull, a local anesthetic (2% Lidocaine hydrochloride; Bela-pharm, Germany) was injected subcutaneously. Further steps were performed under constant and carefully observed Isoflurane anesthesia. All stereotaxic coordinates used are based on the mouse brain atlas (Paxinos and Franklin, 2019).

#### a. Viral microinfusions

All virus suspensions were infused using a 30-gauge (G) needle (Hamilton, USA) through a stereotactic-mounted micropump (UMP3T, World Precision Instruments, UK) at a flow rate of 100 nl/min. After infusion, the needle was left in place for 10 min to ensure adequate virus spread. The drill hole in the skull was closed using bone wax (Ethicon, USA), and the incision was sutured using sterile nylon material and treated with Lidocaine (2% Lidocaine hydrochloride; Bela-pharm, Germany). The following viral constructs (procured from Addgene) were used:

- For Ca^2+^ imaging, a total of 250 nl of AAV-DJ-EF1a-DIO-GCaMP6m (titer of 4 x 10^12^ GC/mL, diluted 1:1.5 in saline; UNC Vector Core) were microinfused in the left caudal LS (LSc; 2 infusions; anterior-posterior (AP): +0.15 mm, medial-lateral (ML): -0.5 mm, and dorsal-ventral (DV): -3.35 mm and - 3.0 mm).
- For chemogenetic experiments, 150 nl of AAV9-hSyn-DIO-hM3Di-mCherry-WPRE (titer 6 x 10^12^ GCs/mL) were microinfused in the left and right LSc (1 infusion/hemisphere; AP: +0.15 mm, ML: +/-0.5 mm, and DV: -2.8 mm).

#### b. Guide cannula implantation

For local infusion, 8-mm-long 23 G guide cannulas were implanted bilaterally 2 mm above the LSc (AP: +0.15 mm, ML: +/-0.5 mm, DV: -1.6 mm) 1 week prior to behavioral testing. The cannulas were fixed to two stainless steel screws using dental cement (Kallocryl, Speiko-Dr. Speier GmbH, Germany). Mice were single-housed after surgery and allowed to recover for at least 5 days. Animals were handled daily to habituate them to infusion procedures and to minimize nonspecific stress responses on the day of the experiment. All guide cannulas were closed with a stylet, which was cleaned daily during handling.

#### c. Grin lens implantation

For Ca^2+^ imaging, the Gradient Refractive Index (GRIN) lens was implanted immediately after viral microinfusion (AP: +0.15 mm, MS: -0.5 mm, and DV: -3.1). ProView^TM^ integrated lenses (Inscopix, USA) were used, with the lens (0.5 NA, 1.5 pitch, 0.5 x 6.1 mm) attached to a baseplate. The craniotomy drilled for the virus was extended for lens implantation using a trephine bit (1.8-mm-diameter tip; Fine Science Tools, USA). The integrated lens was attached to an nVista dummy microscope mounted on a Pro-View stereotax rod (Inscopix, USA) and slowly lowered at 100 µm/min from DV 0 to -3.1 mm. The skull was thoroughly dried, and the lens was secured to the skull with Metabond (Parkell, USA). After the cement had dried, the skin was sutured tightly, and the baseplate was covered with a protective cover (Inscopix, USA).

### Pharmacological manipulations and drug preparations

For pharmacological manipulation of OXT signaling in the LS, the following solutions were bilaterally microinfused into the LS between 30 and 10 min prior to social fear extinction: sterile Ringeŕs solution vehicle control (Veh; B. Braun, Melsungen AG, Germany; 0.2µL/per hemisphere), OXT (0.5ng/0.2µL; Sigma Aldrich: O6379), TGOT (0.5ng/0.2µL; MedChemExpress: Cat#HY-P3467), Atosiban (Ato; 20ng/0.2µL; Tocris: Cat#6332), and Carbetocin (Carb; 0.4µg/0.2µL; Tocris: Cat#4852). All substances were freshly diluted from stock solutions on the day of the experiment. For intra-LSc infusions, the stylets were removed from the left and right guide cannulas and replaced by the infusion cannula (27 G, 10 mm). After infusions, the cannula was left in place for 10 s to ensure adequate local drug diffusion. The animals were carefully observed upon their return to the home cage. Signs of convulsions or tremors were not observed after infusion. For validation of the infusion sites, mice were locally infused with ink postmortem.

### In vivo Ca^2+^ imaging and analysis

#### a. Single-cell microendoscopic calcium imaging

Calcium imaging was performed using nVista (Inscopix, Bruker) miniscopes attached to the previously implanted baseplate after removal of the protective cover. Correct positioning of the miniscope was achieved via the built-in magnetic coupling, and focus was adjusted individually for each animal before recording. Signals were acquired at 10 Hz with a mean exposure time of 100 ms using Inscopix Data Acquisition Software (IDAS). Excitation light intensity was optimized for each session based on fluorescence signal quality and typically ranged from 0.8 to 1.2 mW/mm². Temporal alignment between calcium imaging and behavioral tracking was achieved using TTL signals triggered from the nVista DAQ-box. Behavior was monitored with monochrome cameras operating at 25 frames/s. Raw imaging data were preprocessed in Inscopix Data Processing Software (IDPS), including spatial downsampling, spatial bandpass filtering, and motion correction. Identification of cellular components and extraction of fluorescence traces were performed using a custom MATLAB pipeline (MathWorks, R2017b) based on the CNMF-E framework ^25^, and were subsequently inspected manually. For analyses across repeated recordings, longitudinal cell matching was carried out using automated registration with the MATLAB-based CellReg GUI ^26^ followed by manual curation.

#### b. Population-level comparison of normalized calcium activity

Calcium imaging data were analysed in relation to annotated behavioral states in the SFC paradigm. For population-level analyses, calcium activity was normalized to baseline periods yielding baseline-normalized z-scored activity traces. Baseline was defined as the 90 s period immediately preceding and following each social or non-social stimulus presentation. The social approach, contact, and retreat were pooled into a single “social” category, whereas all remaining annotated behaviors were grouped as “other”.

#### c. Classification of stimulus-responsive cells

To identify neurons modulated around social contact, behavioral annotations were transformed into binary predictors indicating (i) direct social contact, (ii) the 3s period preceding contact onset, and (iii) the 3s period following contact offset. For each neuron, separate linear models were fit in R using the lm function. Resulting p-values were corrected for multiple comparisons across neurons for each predictor term using the Hochberg method. Neurons were classified as Excited or Inhibited according to the sign of the regression coefficient when the adjusted p-value was < 0.05, and as non-responsive otherwise.

#### d. Identification of stimulus response profile

To quantify calcium dynamics around behavioral state transitions, baseline-normalized calcium traces were segmented into discrete behavioral episodes defined as contiguous runs of the same annotated behavior. Episodes shorter than 1 s were excluded from further analysis. For each valid episode, peri-event traces were extracted for behavioral entries in windows spanning 5 s before to 5 s after the event. Episodes were included only when the complete peri-event window was available; events for which the window extended beyond the beginning or end of the recording were excluded.. Baseline activity was defined as the 5 s period immediately preceding episode onset. In each case, the raw calcium signal was converted to a z-scored peri-event trace relative to the mean and standard deviation of the corresponding baseline window.

#### e. Construction of neural features and neuronal score

During the extinction phase, candidate neural features were derived from both calcium activity and linear-model-based contact modulation estimates ^27, 28^. Calcium traces from the social and non-social stimulus conditions were used to compute single-cell signal features, including mean activity during pre-contact, contact, and post-contact periods, as well as derived contrasts such as contact minus pre-contact, post-contact minus pre-contact, and social minus non-social contact activity. These single-cell features were then aggregated per animal by calculating the mean and median across cells, and the number of contributing cells was retained for each feature. In parallel, linear-model outputs from the extinction dataset were used to derive per-animal modulation features for social and non-social contact, including average contact beta values, the fraction of modulated, excited, and inhibited cells, and a net modulation index defined as the fraction of excited minus inhibited cells. Three social-contact-related features - social post-minus-pre activity, mean social contact activity, and the mean social contact beta coefficient - were selected for score construction. For each selected feature, per-animal values were z-scored across animals and averaged to generate a neuronal score for each mouse.

### Chemogenetic manipulation

For chemogenetic manipulation of LS^OXTR^, AAV9-hSyn-DIO-hM3Di-mCherry-WPRE was injected in LS of OXTR-CRE mice (1 infusion/hemisphere; AP: +0.15 mm, ML: +/-0.5 mm, and DV: -2.8 mm), which led to the Cre-driven expression of an inhibitory designer receptor activated by designer drug (Gi-DREADD)., which led to the Cre-driven expression of an inhibitory designer receptor activated by designer drug (Gi-DREADD). Behavioral testing was performed 3 weeks after injection. Mice received an intraperitoneal (i.p.) infusion of either clozapine-n-oxide (CNO; 10 mg/kg in 0.2 ml of 0.9% saline) or Veh (0.2 ml of 0.9% saline) 30 min prior to social fear extinction.

### Neuroanatomical analysis

#### a. Receptor autoradiography

All brains were removed, snap frozen, and stored at −20 °C. OXTR autoradiography was performed on 16-μm coronal sections targeting the LS (Bregma 1.1 to −0.1 mm) as previously described using a linear OXTR antagonist [^125^I]-d(CH_2_)_5_[Tyr(Me)_2_-Tyr-Nh_2_]_9_-OVT (Perkin Elmer) ^10^. The exposure was set to 5 days based on OXTR density in the region of interest. OXTR binding was analyzed using ImageJ (V1.37i, National Institutes of Health) and calculated for each mouse by averaging 3 sections per region of interest. The background signal was subtracted to control for non-specific binding. Left and right regions were scored separately and pooled if no significant hemispheric difference was found.

#### b. Immunohistochemistry

To study the distribution pattern of LS^OXTR^, male OXTR-Cre:Ai9 mice were transcardially perfused with paraformaldehyde (PFA) under deep anesthesia 90 min after social fear acquisition. Brains were removed and post-fixed in 4% PFA for 24 h, followed by cryo-protection in 30% sucrose for 2 days and snap-freezing in isopentane. Forty-µm coronal cryocut slices (CM3000, Leica, Germany) containing the LS (Bregma from +1.18 to - 0.15) were kept in PBS containing 5% normal goat serum (Vector Laboratories) and 0.5% TritonX-100 (1 h at room temperature (RT), and afterward, incubated with primary anti-OXT antibody (PS38; provided by Dr. Harold Gainer). Appropriate secondary antibody (Anti-rabbit IgG, 7074S; Cell Signaling Technology) was applied (2 h, RT) in the dark. Slides were covered with SuperFrost slides using Roti® Mount FluorCare DAPI (Carl Roth, Germany). Analysis of density was performed by counting the number of LS^OXTR^ within the rostral (LSr), LSc using ImageJ (V1.37i, National Institutes of Health).

#### c. RNAscope

Visualization of single RNA molecules was carried out using RNAscope. This in situ hybridization technique was performed according to the manufactureŕs protocol (RNAscope Multiplex Fluorescent V2 Assay, Advanced Cell Diagnostics). Briefly, fresh frozen brain slices were stored at -80°C until further processing. Sections were post-fixed with 4 % PFA in 0.1 M PBS cooled to 4°C for 30 min. Sections were then dehydrated in 50% and 70% ethanol, and twice in 100% ethanol. After drying, hydrogen peroxidase was applied for 10 min in order to remove blood residues. Cells were then permeabilized by 10 min Protease III treatment and washed with 0.1 M PBS. Tissue was then covered with RNAscope probes targeting OXTR (412171-C1), Somatostatin (SSt; 404631-C3), and Neurotensin (Nts; 420441-C3) at 40°C for 2 h. After probe hybridization, sections were washed twice for 2 min in RNAscope Wash buffer, then sequentially incubated with RNAscope Amp-I and Amp-II for 30 min and Amp-III for 15 min, with two washes in Wash buffer between each treatment. Signals were developed using the TSA Plus fluorophore Cy5 (1:1000), Cy3 (1:1000) and Flp (1:1000) (FP1168, FP1170, FP1171, perkin Elmer). Sections were covered with Roti-Mount FlurCare DAPI. Images were obtained with a confocal microscope (name, company, country) at 40x magnification at different subregions of the LS. Images were analyzed using a custom Cell Profiler Pipeline ^29^.

### Statistical analysis

All data were analyzed using SPSS (Version 26.0, IBM Corp., USA), GraphPad Prism (Version 8, GraphPad Software, San Diego, USA), or R (v4.3.3). All datasets were first tested for normal distribution using SPSS (Version 26.0, IBM Corp., USA) or R (v4.3.3). Accordingly, parametric paired and unpaired t-tests were performed between two groups, or non-parametric Wilcoxon and Mann-Whitney U-tests were used when the data did not meet the assumptions of normality. Analysis of variance (ANOVA) with a mixed model was used to assess two factors across two or more time points (factor sex; factor treatment x SFC x time, in SFC extinction and recall). The Kruskal-Wallis test or the Geisser-Greenhouse correction was applied when sphericity was violated. Statistical significance was accepted at p ≤ 0.05, while a trend was recognized at p ≤ 0.07. Data is represented as the mean ± standard error of the mean (SEM).

## Results

### A caudally enriched population of LS^OXTR^ neurons is recruited by aversive social stimuli

Given the substantial molecular and anatomical heterogeneity of the LS, we first characterized the precise anatomical distribution of LS^OXTR^ neurons. Using receptor autoradiography, we observed significantly higher OXTR binding in the LSc (Bregma: 0.5 to -0.1) in comparison to the rostral LS (LSr; Bregma: 1.1-0.5) (Fig. 1b). The rostro-caudal increase in OXTR binding was corroborated by the finding of a larger population of OXTR^+^ neurons in the LSc (LSc^OXTR^) of OXTR-Cre:*Ai9* mice (Fig. 1c). Neurotensin (Nts) and somatostatin (SSt), which also regulate behavioral responses to social trauma ^30, 31^, are well-established neuropeptidergic markers for LS GABAergic neurons. Hence, we further quantified co-expression of Nts and SSt in LSc^OXTR^ neurons using RNAScope. While 21.4% of LS cells express OXTR, the majority (60.2%) of OXTR^+^ cells lack co-expression of Nts or SSt (Fig. 1d), and only smaller fractions co-express Nts (8.3%), SSt (15.5%), or both (16.1%). These data show a pronounced enrichment of LS^OXTR^ neurons in the LSc, most of which represent a distinct molecular subtype of LS GABAergic neurons. We next used immunohistochemistry to reveal that OXT^+^ fibers are distributed throughout the LS, with no differences between LSr and LSc (Fig. 1e), suggesting that the LS is uniformly innervated by hypothalamic OXTergic projections.

**Fig. 1:**
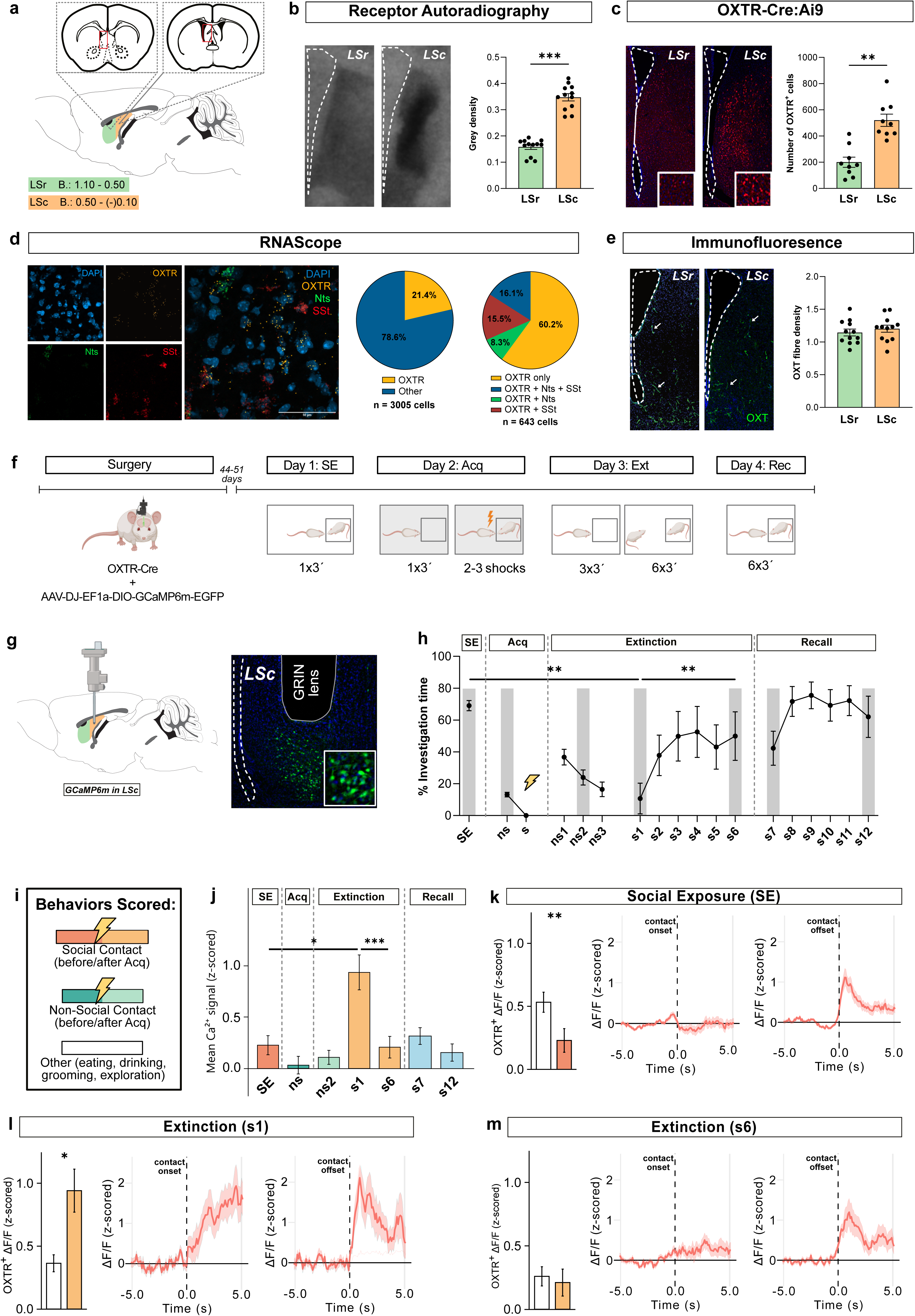
Distribution of oxytocin (OXT) receptor-expressing cells (OXTR^+^) in the lateral septum (LS) and their activity during exposure to social and non-social stimuli before and after social fear conditioning (SFC). **a**, Schematic representation of brain slices from the rostral (LSr) and caudal (LSc) LS showing the area (red outline) represented in images in panels b, c, and e. b, LS brain sections with radioactive-labeled OXTR and average grey density in the LSr (green) and LSc (orange) (n = 12 images (4 mice)). b, LS brain section from OXTR-Cre:Ai9 mice showing OXTR^+^ neurons (red) and the total number of OXTR^+^ neurons in LSr and LSc (n = 9 mice). c, Triple in situ hybridization showing colocalization of Oxtr with neurotensin (Nts) and somatostatin (SSt), and the percentage of Oxtr^+^ cells in the LS (left) and co-expression of Oxtr with Nts and SSt (right) (n = 4 mice). e, LS brain sections showing immunostained OXT fibers and average OXT fiber density in the LSr (green) and LSc (orange) (n = 12 images (3 images/mouse)). f, Schematic of the experimental timeline for miniscope-based Ca^2+^ imaging during the adapted SFC paradigm (created using BioRender.com). g, GRIN lens placement scheme (created using BioRender.com) and GCaMP6s expression in the LSc. h, Percentage time mice spent investigating non-social (ns) and social (s) stimuli in the adapted SFC paradigm (SE: Social exposure; Acq: Acquisition; Ext: Extinction; Rec: Recall; n = 9 mice). Grey bars represent sessions during which Ca^2+^ activity was measured. i, Description of the behaviors assessed during various sessions of the adapted SFC paradigm. j, Ca^2+^ activity measured during social investigation within the sessions indicated by grey bars in h (SE: n = 39, ns: n =73, ns2: n = 90, s1: n = 78, s6: n = 66, s7: n = 108, s12 = 117). k-m, Comparison of Ca^2+^ activity during social investigation with other behaviors and average Ca^2+^ activity traces during SE (k; n = 39 for social, n = 39 for other), s1 (l; n = 78 for social, n = 106 for other), and s6 (m; n = 66 for social, n = 106 for other). Data represents mean grey density ± S.E.M (b), mean number of OXTR^+^ neurons ± S.E.M (c), mean OXT fiber density ± S.E.M (e), mean percentage investigation ± S.E.M (h), and mean ΔF/F ± S.E.M (j, k, l). *p≤0.05 (j, k), **p≤0.01 (c, k), ***p≤0.001 (b, j).

To monitor the real-time activity of LSc^OXTR^ neurons during social interaction before and after social trauma (i.e., social fear acquisition), we combined an adapted SFC paradigm with Ca^2+^ imaging using miniscopes. For this purpose, we used Cre-driven expression of a Ca^2+^ indicator (GCaMP6m) in LSc^OXTR^ neurons (Fig. 1f, g). One day prior to social fear acquisition, mice were exposed to an age- and sex-matched conspecific in their home cage to assess baseline social behavior (day 1). During social exposure (SE), they spent approximately 70% of time interacting with the conspecific, indicating high sociability (Fig. 1h). As a result of social fear acquisition performed on day 2, we observed a significant reduction in social investigation during exposure to the first social stimulus (s1) of extinction training on day 3, (s1 vs SE; Fig. 1h). However, during exposure to s6, i.e., after consecutive exposure to 5 different social stimuli, sociability was restored to pre-acquisition levels i.e., during SE on day 1 (approximately 70%) (Fig. 1h). During recall (s7-s12; day 4) mice exhibited high social investigation indicative of robust extinction of social fear. Importantly, the time spent investigating non-social objects (ns1-ns3) did not differ across sessions, demonstrating that social fear was specifically directed toward social stimuli and did not alter overall investigatory behavior (Fig. 1h).

Measurement of social investigation-associated Ca^2+^ dynamics during social exposure (SE; day 1), extinction (Ext; s1 and s6), and recall (Rec; s7 and s12) revealed that the activity of LSc^OXTR^ neurons was closely linked to the valence of the social stimulus. LSc^OXTR^ neurons displayed low activity during social interaction within the SE session, when the social stimulus had a positive valence (day 1; Fig. 1j, k). In contrast, their activity was prominently and selectively enhanced upon social investigation during s1 (i.e., the start of social fear extinction) following acquisition-induced association of the social stimulus to a negative valence (day 3; Fig. 1j, l). Interestingly, recovery of approach behavior at the end of extinction (Fig. 1h) was associated with reduced social investigation-associated activity of LSc^OXTR^ neurons (s6; Fig. 1j, m), comparable to social interaction prior to social fear acquisition (Fig. 1k). LSc^OXTR^ neurons remained largely inactive during the investigation of non-social stimuli (ns and ns2; Fig. 1j) and during display of non-social behaviors like feeding, drinking, grooming, and general exploration (Fig. 1k-m). These data indicate that LSc^OXTR^ neurons are specifically recruited during interactions with aversive social stimuli.

### Valence-dependent LSc^OXTR^ dynamics track extinction success

We hypothesized that individual variability in social interaction across different sessions of the SFC paradigm might be reflected by distinct activity dynamics of LSc^OXTR^ neurons. To test this, we retrospectively classified mice from the prior cohort (Fig. 1h) into those that did not respond to extinction training (NRes; Fig. a, b, d) and still expressed social fear at its end and those that responded to social fear extinction training (Res; Fig. a, c, e) with a reinstatement of sociality, as previously described ^7^. During general social exposure (SE; day 1) and the first social stimulus of extinction (s1; day 3), both NRes and Res mice exhibited similarly high and low levels of social investigation, respectively, suggesting that social fear acquisition successfully induced social avoidance in both groups (Fig. 2a-e). However, following repeated social exposure (s1-s6) during extinction, NRes mice did not respond to extinction, as reflected by continued low levels of social investigation and reduced number of social investigation bouts during s6, which were comparable to s1 and significantly lower than those of Res mice (Fig. 2a, b, d). In contrast, Res mice recovered from acquisition-induced social fear, reflected by high levels of overall social investigation and the number of social investigation bouts during the final extinction session (s6; Fig. 2a, c, e). Comparison of the activity of LSc^OXTR^ neurons of NRes (Fig. 2f) and Res (Fig. 2g) mice during SE (day 1) and s1 (day 3) revealed low activity during social investigation in the SE session and elevated activity during social investigation during s1 exposure in both cohorts. Strikingly, at the end of extinction (s6), a marked social investigation-associated inhibition of LSc^OXTR^ neurons (similar to the SE session) was observed only in Res mice (Fig. 2d, e). Thus, extinction-induced inhibition of LSc^OXTR^ neurons is associated with extinction success.

**Fig. 2:**
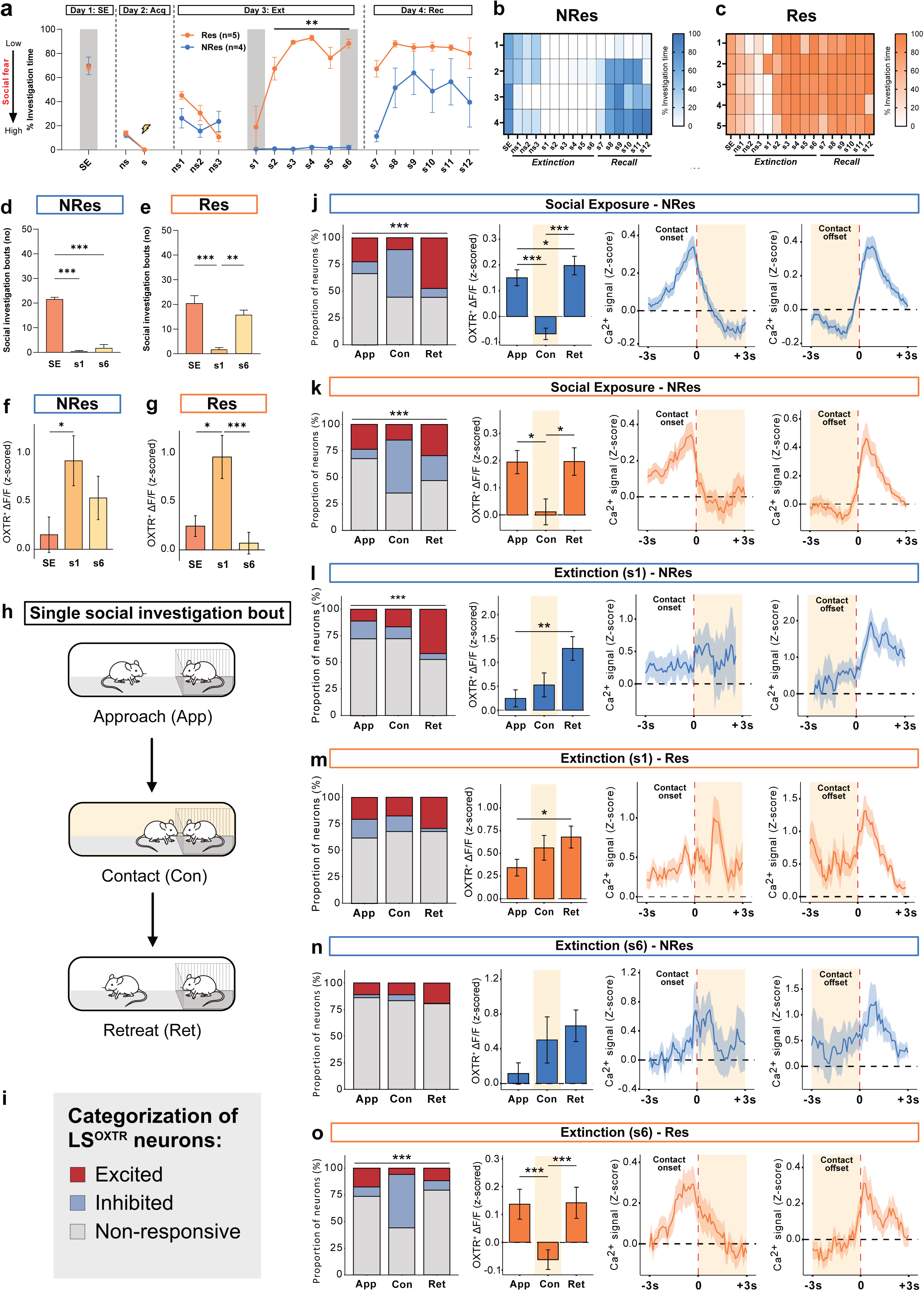
Activity dynamics of oxytocin receptor (OXTR) expressing neurons in the caudal lateral septum (LSc^OXTR^) during various phases of the social fear conditioning (SFC) paradigm in non-responder (NRes; Blue) and responder (Res; Orange) mice. **a**, Percentage of time mice spent investigating non-social (ns) and social (s) stimuli in the SFC paradigm (SE: Social exposure; Acq: Acquisition; Ext: Extinction; Rec: Recall) in NRes and Res mice. **b, c,** Heatmaps showing the percentage of time spent with social investigation of NRes (b) and Res (c) mice during SE, Ext, and Rec. **d, e,** Graphs showing average number of social investigation bouts in NRes(b) and Res (c) mice. **f, g,** Ca^2+^ activity of NRes (f; n = 32 for SE, n = 38 for s1, n = 46 for s6) and Res (g; n = 47 for SE, n = 40 for s1, n = 20 for s6) mice measured during social investigation within SE, s1, and s6 sessions. **h,** Schematic representation of the temporal stratification of a single social investigation bout into approach (App), contact (Con), and retreat (Ret) phases. **i,** Legend for the categorization of LSc^OXTR^ neurons into those that are excited (red), inhibited (light blue), or non-responsive (grey). **j-o,** Percentage of excited, inhibited, and non-responsive LSc^OXTR^ neurons of NRes (j, l, n; n = 59, 60, and 60 neurons, respectively) and Res (k, m, o; n = 40, 46, and 46 neurons, respectively) mice along with a representative neuronal trace during SE (Day 1; j, k), s1 (Day 3; l, m), and s6 (Day 3; n, o) sessions. Data represent mean percentage investigation ± S.E.M (a), average number of social investigation bouts (b, c), mean ΔF/F ± S.E.M (f, g), percentage of neurons (panel 1), mean OXTR ΔF/F ± S.E.M (panel 2), and Ca^2+^ signal Z-score for contact onset (panel 3) and offset (panel 4) (j-o). *p≤0.05 (d, e), **p≤0.01 (a, c, e), ***p≤0.001 (b, c, g, j, k, l, o).

Social behaviors are intrinsically dynamic, raising the possibility that different phases of a social investigation bout are encoded by distinct patterns of neuronal activity. Such patterns may flexibly reorganize in SFC^+^ mice as the valence of a social stimulus is updated during extinction. For this purpose, each bout of social investigation was divided into approach, contact, and retreat phases (Fig. 2h). We then analyzed the activity of LSc^OXTR^ neurons during these phases within SE (day 1), s1 (extinction, day 3), and s6 (extinction, day 3) sessions. In the SE session, the majority of LSc^OXTR^ neurons in both NRes (Fig. 2j) and Res (Fig. 2k) mice were inhibited upon contact with the conspecific. This contact-associated inhibition of LSc^OXTR^ neurons was absent in both NRes (Fig. 2l) and Res (Fig. 2m) cohorts during s1, i.e., at the start of extinction when the social stimulus is aversive (i.e., has a negative valence). However, during s6 (i.e., the end of extinction), the Res mice exhibited neuronal dynamics similar to SE, with inhibition of the majority of LSc^OXTR^ neurons during contact with the conspecific (Fig. 2o), whereas, in contrast, no inhibition of LSc^OXTR^ neurons was observed in NRes mice (Fig. 2n). Interestingly, by the end of recall session (i.e., s12), when the NRes mice no longer exhibited social fear (Supplementary Fig. 1a, b), the contact-associated inhibition of LSc^OXTR^ neurons that returned to similar levels as in the SE session (Supplementary Fig. 1f, h) and in Res mice (Supplementary Fig. 1g, i). These data suggest that inhibition of LSc^OXTR^ neurons specifically during direct social contact is associated with positive social valence. Thus, restoring the modulatory pattern of LSc^OXTR^ neurons observed during the SE session might drive the extinction-associated reassignment of positive valence to social stimuli.

### LSc^OXTR^ neuronal activity distinguishes extinction success

To determine whether extinction-related neuronal activity could be integrated into a single phenotype-relevant measure, we constructed a composite neuronal score from three complementary per-mouse features derived from extinction sessions: mean activity during social contact, post-contact activity increase, and the social contact modulation coefficient (Fig. 3a). Mean activity during social contact (Fig. 3c) directly quantifies the net excitation or inhibition of LSc^OXTR^ neurons during the critical phase of social contact within each investigation bout (Fig 2h). The post-contact activity increase (Fig. 3d) quantifies the recovery of neuronal activity after social contact offset, defined as the difference in Ca^2+^ signal between the 3-s post-contact and 3-s pre-contact windows. The social contact modulation coefficient (Fig. 3e) provides a linear model-based estimate of the extent to which neuronal activity is modulated during social contact.

**Fig. 3:**
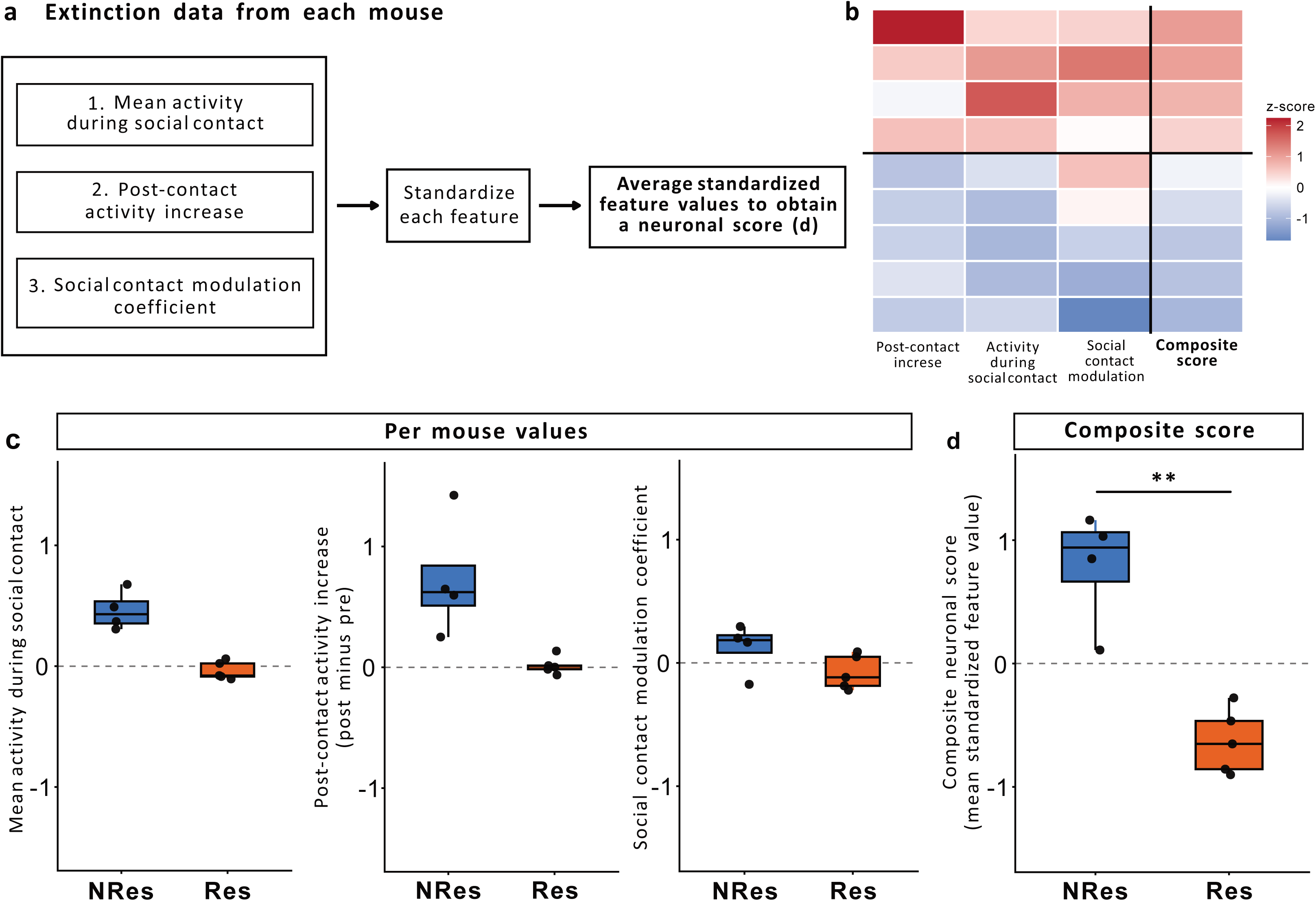
Activity of oxytocin receptor (OXTR)-expressing neurons in the caudal lateral septum (LSc^OXTR^) distinguishes extinction success in social fear conditioning. **a**, Schematic illustrating the construction of the neuronal score from per-mouse features. Three neural features were extracted from each mouse: post-contact activity increase, mean activity during social contact, and the social contact modulation coefficient. Each feature was standardized across mice and then averaged to generate a composite neuronal score. **b**, Heatmap showing the standardized feature profile of individual mice. Rows represent individual mice and columns represent the three standardized component features together with the composite neuronal score. **c**, Per-mouse values of the three component features in NRes and Res mice. **d**, Composite neuronal score in NRes and Res mice. The neuronal score significantly differed between phenotypes (Wilcoxon test, p = 0.02).

Each feature was standardized across mice to obtain a single neuronal score for each animal. Inspection of the standardized feature profiles revealed a clear phenotypic organization, with NRes mice showing predominantly positive values across all three component features – higher activity during social contact (Fig. 3c), greater post-contact activity increase (Fig. 3d), and a more positive contact modulation coefficient (Fig. 3e), whereas Res mice exhibited predominantly lower or negative values. Integrating these convergent neuronal measures into a single score for each mouse yielded a distinct separation between phenotypes, with NRes mice displaying positive scores and Res mice displaying negative scores (Fig. 3d). These data indicate that extinction-associated neuronal dynamics are organized along a continuous axis that distinguishes NRes and Res mice and can be captured by a simple composite neuronal score. Importantly, this score suggests that extinction success is better explained by the combined temporal dynamics of LSc^OXTR^ neurons before, during, and after social contact than by any single feature alone. Thus, LSc^OXTR^ neuronal activity is closely associated with social valence updating during extinction, with transient inhibition during social contact emerging as a neural signature of successful extinction.

### OXTergic fibers differentially innervate the LS in Res and NRes mice

The LS receives topographically organized inputs from various brain structures ^8, 9^, including OXTergic projections from the SON, which are essential for social fear extinction ^6^. Moreover, we have also shown that OXT release in the LS is associated with enhanced social investigation ^6, 10^. However, the pattern of OXTergic innervation in the LS that may underlie differential signaling and consequent differences in the extinction success of Res and NRes mice remains unknown. Therefore, we subjected OXTR-Cre:*Ai9* mice to the SFC paradigm and characterized OXTergic fibers within the LS of Res and NRes mice. As before, mice were retrospectively classified as Res or NRes based on their behavior during social fear extinction (Fig. 4b, c). Despite receiving the same number of CS-US pairings during acquisition (Fig. 4a), Res mice responded to the extinction training with enhanced social contact (above 45% of total time) during exposure to the last two social stimuli (s5-s6; Fig. 4b, c). In contrast, in NRes mice, social investigation remained low (below 45% of total time) during s5 and s6 exposure as previously described ^7^. OXTergic fibers in the LSc were located in close proximity to LSc^OXTR^ neurons (Fig. 4e). While the OXTergic innervation within LSr (Fig. 4e, h) and LSc (Fig. 4f, i) did not differ between NRes and Res mice, significantly more OXTergic fibers were found towards the LSc (versus LSr) in Res mice than NRes mice (Fig. 4j). These data, in combination with our previous results ^6, 10^, suggest that increased OXT signaling in the LSc contributes to the success of social fear extinction in Res mice.

**Fig. 4:**
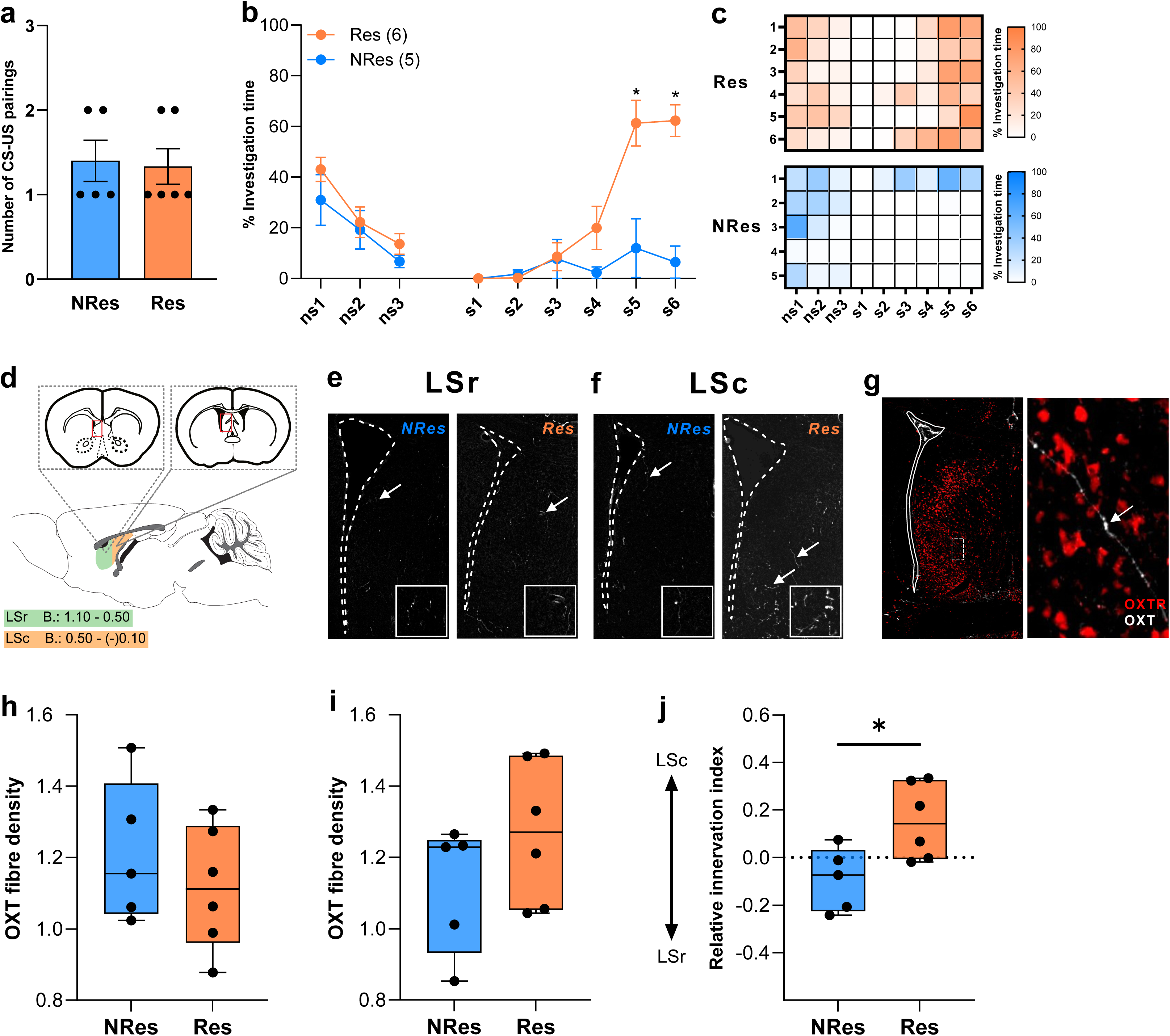
Distribution of oxytocin (OXT) -positive fibers in the lateral septum (LS) of non-responder (NRes) and responder (Res) mice. **a**, Number of conditioned stimulus (CS) -unconditioned stimulus (US) pairings received during social fear acquisition (day 1). **b**, Percentage time spent investigating the non-social (ns1-ns3) and social (s1 - s6) stimuli during social fear extinction (day 2). **c**, Heatmap showing the percentage of time spent on social investigation of Res (n=6) and NRes (n=5) mice during social fear extinction. **d**, Schematic representation of the rostral (LSr) and caudal (LSc) LS in sagittal and horizontal brain slices. **e, f**, Representative immunofluorescence images from the LSr (e) and LSc (f) of NRes and Res mice. **g**, Representative immunofluorescence image from the LSc showing the OXT fiber (white) in proximity to LSc^OXTR^ neurons (red). **h**, **i**, OXT fiber density within the LSr (h) and LSc (j) of NRes and Res mice. **j**, Relative innervation index for OXT fibers within the LS. Data represents mean CS-US pairings ± S.E.M (a), mean percentage investigation time ± S.E.M (b), OXT fiber density ± S.E.M (h, i), and relative innervation index ± S.E.M (j). *p≤0.05 (b, j).

### LSc^OXTR^ neurons are essential for the successful extinction of social fear

Building on the finding that the temporally precise modulation of LSc^OXTR^ neurons encodes successful extinction, we investigated their causal role in regulating social fear extinction using chemogenetic inhibition (Fig. 5a-g). For chemogenetic inhibition, OXTR-Cre mice received bilateral injections of a Cre-dependent Gi-DREADD into the LSc 3 weeks prior to exposure to the SFC paradigm, generating groups of SFC^+^ and SFC^-^ mice that received a pre-extinction ip injection of either CNO or vehicle. During social fear acquisition, a comparable number of CS-US pairings was required to induce avoidance of the conspecific in both SFC^+^/CNO and SFC^+^/Veh groups prior to ip treatment (Fig. 5c). On the following day, mice received either CNO (10mg/kg, ip) or vehicle (Veh) 30 min prior to social fear extinction. Evidencing successful social fear conditioning, both SFC^+^/CNO and SFC^+^/Veh groups avoided the first social stimulus, whereas their respective SFC^-^ controls displayed high levels of social investigation (s1; Fig. 5d). The SFC^+^/Veh mice underwent successful extinction reflected by rising social investigation times until they achieved the level of social contact displayed by the SFC^-^/Veh group (s6; Fig. 5d). In contrast, chemogenetic inhibition of LSc^OXTR^ neurons in SFC^+^/CNO mice prevented social fear extinction reflected by low levels of social investigation still seen at the end of social fear extinction (s6; Fig. 5d). Pre-extinction infusion of CNO did not influence social behavior in SFC^+^ mice in comparison to Veh-injected controls (Supplementary Fig. 2), suggesting the state-specific role of LSc^OXTR^ neurons in regulating extinction-associated valence reasignment. Interestingly, only SFC^+^/CNO mice still expressed social fear during the beginning of recall (day 3), reflected by reduced social investigation compared to all other groups (s7-s9; Fig. 5d), which was only reversed at the end of recall (s10-s12; Fig. 5d). Inhibition of LSc^OXTR^ neurons significantly reduced the proportion of Res mice within the SFC^+^ groups (Fig. 5e, f, g) further supporting the necessity of LSc^OXTR^ neurons for the adaptive social learning essential for successful social fear extinction. Together, these data demonstrate that LSc^OXTR^ neurons, and specifically their precise temporal activity patterns, are necessary for updating social valence during extinction, enabling the transition from a negative to a positive social representation and, consequently, recovery from acquisition-induced social fear.

**Fig. 5:**
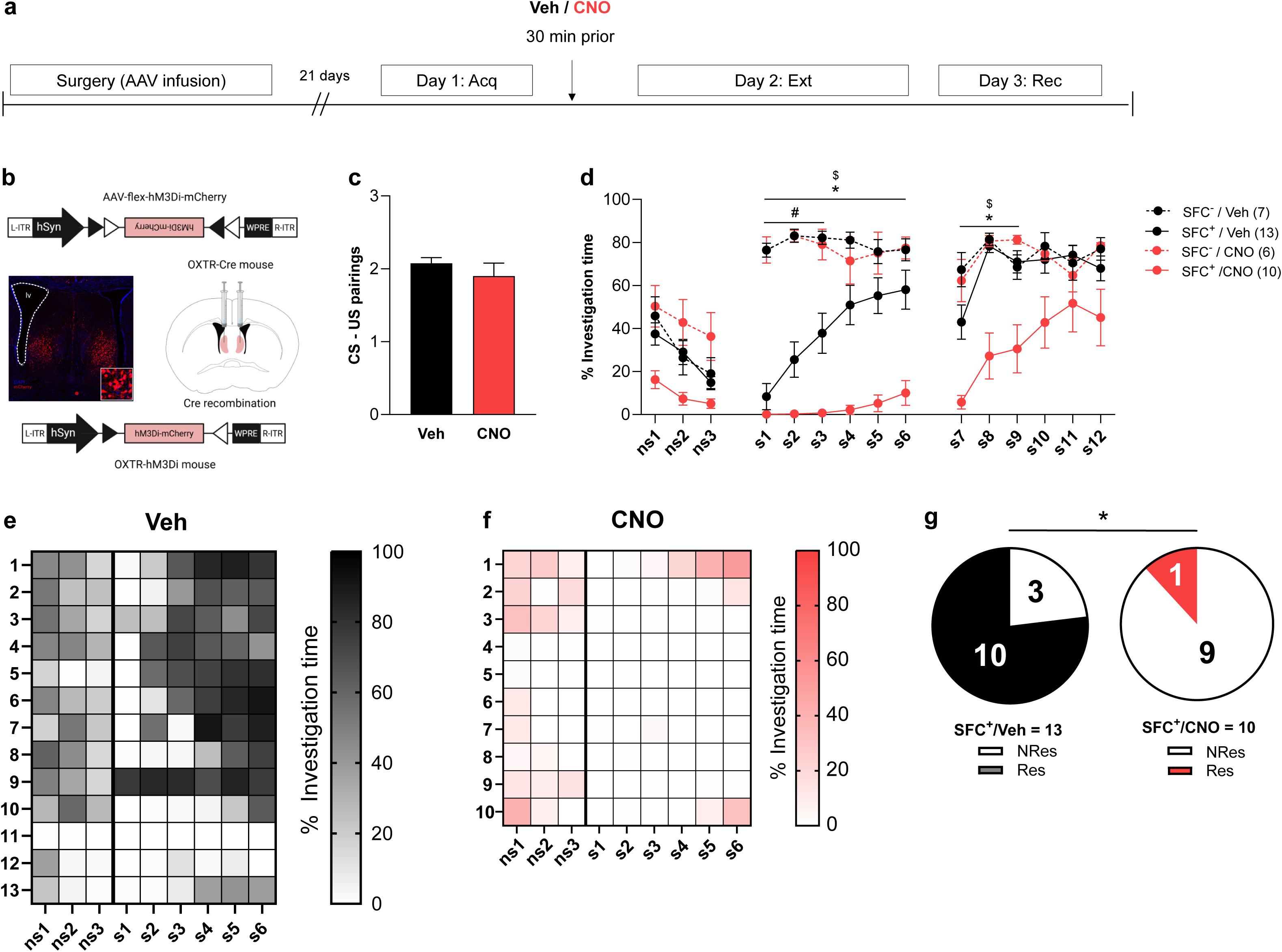
Effect of chemogenetic inhibition of oxytocin receptor (OXTR) expressing neurons in the caudal lateral septum (LSc^OXTR^). **a**, Timeline for the chemogenetic experiment including surgery (bilateral microinfusions of AAV9-hSyn-DIO-hM3Di-mCherry-WPRE (150nl/hemisphere) in the LSc of OXTR-Cre mice), and social fear conditioning (SFC) paradigm (Day 1-3). **b,** Sketch of the infusion scheme along with a representative image of viral expression for chemogenetics**. c**, Number of conditioned-unconditioned (CS-US) stimulus pairings received by social fear conditioned (SFC^+^) mice during social fear acquisition (Acq). **d**, Social investigation times during social fear extinction (ns1-s6; Ext) and recall (s7-s12; Rec) of SFC^+^ and unconditioned (SFC^-^) mice treated with either clozapine-n-oxide (CNO; 10 mg/kg/0.2ml of saline; i.p.) or vehicle (Veh; 0.2ml of saline; i.p.) 30 min prior to extinction. **e, f,** Heatmaps showing the percentage of social investigation of SFC^+^/Veh (e) and SFC^+^/CNO (f) mice during Ext. **g**, Pie chart representing the number of non-responder (NRes) and responder (Res) mice within SFC^+^/Veh and SFC^+^/CNO groups. Data represent mean CS-US pairings ± S.E.M (c), mean percentage investigation time ± S.E.M (d), and absolute number of NRes and Res mice (g). *p≤0.05 (d, g; SFC^+^/Veh vs. SFC^+^/CNO), # p≤0.05 (d; SFC^-^/Veh vs. SFC^+^/Veh), $ p≤0.05 (d; SFC^-^/CNO vs. SFC^+^/CNO).

### OXTR-mediated signaling modulates social valence during extinction

To further determine whether OXT signaling via OXTR drives extinction success, we pharmacologically targeted OXTRs in the LSc 10 min prior to extinction training (Fig. 6a). First, we assessed the dose-dependent effects of synthetic OXT bilaterally injected into LSc via previously implanted guide cannulas on social fear extinction (Fig. 6b). During social fear acquisition (day 1), an experimental and a control group of SFC^+^ mice received the same number of CS-US pairings (Fig. 6c). On day 2, 10 min after pre-extinction LS infusion, all groups exhibited similar exploration of non-social stimuli (ns1-ns3; Fig. 6d). Importantly, SFC^+^/OXT mice displayed an accelerated extinction of social fear, as they showed higher levels of social investigation in comparison to SFC^+^/Veh mice during the initial stage of social fear extinction (s2-s3; Fig. 6d). By the end of extinction all groups of mice achieved social investigation levels similar to the SFC^-^/Veh controls (s6; Fig.6d). During recall (day 3), all animals exhibited comparable levels of social investigation (Fig. 6d). Interestingly, only the administration of OXT significantly enhanced the proportion of Res mice (Fig. 6e-g) suggesting that augmentation of OXT signaling is important for efficient re-association of a positive emotional valence to a social stimulus.

**Fig. 6:**
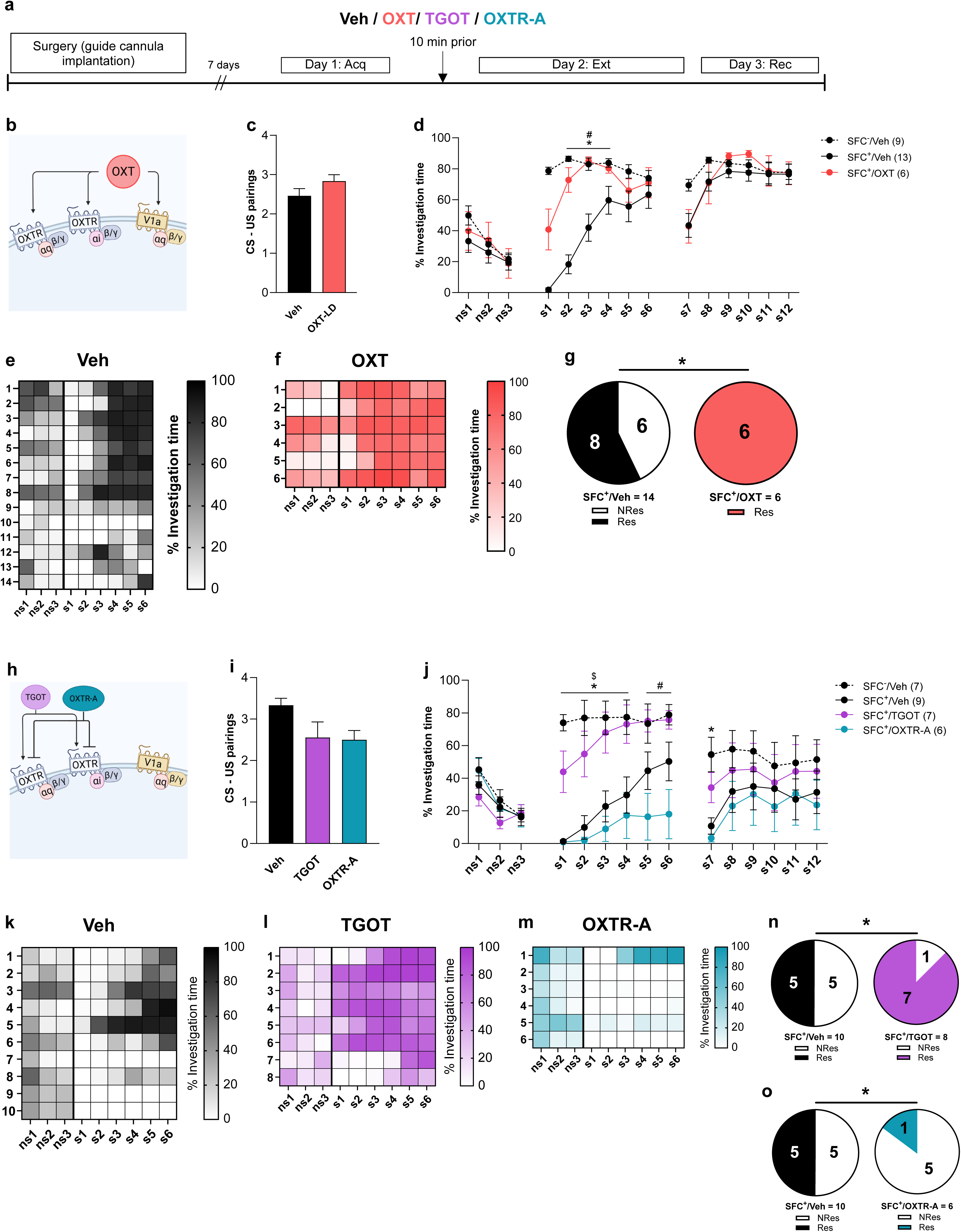
Effect of pharmacological activation or inhibition of oxytocin (OXT) receptors (OXTRs) in the caudal lateral septum (LSc) on social fear extinction. **a**, Experimental timeline. **b**, Scheme of OXT binding to both OXTR and vasopressin V1A receptor. **c**, Number of conditioned – unconditioned stimulus (CS-US) pairings received by social fear conditioned (SFC^+^) mice during social fear acquisition (Acq). **d**, Social investigation times during social fear extinction (ns1-s6; Ext) and recall (s7-s12; Rec) from SFC^+^ and unconditioned (SFC^-^) mice after bilateral infusions of synthetic OXT (0.5ng/0.2µl/side) or vehicle (Veh; Ringer; 0.2µl/side) 10 min prior to Ext. **e**, **f**, Heatmaps showing the percentage of social investigation during Ext in SFC^+^/Veh (e) and SFC^+^/OXT (f) mice. **g,** Pie chart representing the number of non-responder (NRes) and responder (Res) mice in SFC^+^/Veh vs. SFC^+^/OXT mice. **h**, Scheme of TGOT (selective and complete OXTR agonist) and the selective OXTR antagonist (OXTR-A) that exclusively binds the OXTR. **i**, Number of CS-US pairings received by SFC^+^ mice during Acq. **j**, Social investigation times during Ext (ns1-s6) and Rec (s7-s12) from SFC^+^ and SFC^-^ mice that received bilateral infusions of TGOT (5ng/0.2µl/side), OXTR-A (0.5ng/0.2µl/side), or Veh (Ringer; 0.2µl/side) 10 min prior to Ext. **k-m,** Heatmaps showing the percentage of social investigation during Ext in SFC^+^/Veh (m), SFC^+^/TGOT (n), and SFC^+^/OXTR-A (p) mice**. n**, **o**, Pie charts representing the number of NRes and Res mice in SFC^+^/Veh vs. SFC^+^/TGOT (n) and in SFC^+^/Veh vs. SFC^+^/OXTR-A (o) mice. Data represent mean CS-US pairings ± S.E.M (c, i), mean percentage investigation time ± S.E.M (d, j), and absolute number or NRes and Res mice (g, n, o). *p≤0.05 (d: SFC^+^/Veh vs. SFC^+^/OXT, j: SFC^+^/Veh vs. SFC^+^/TGOT, g, n, o), #p≤0.05 (d: SFC^-^ /Veh vs. SFC^+^/Veh, j: SFC^+^/Veh vs. SFC^+^/OXTR-A), $p≤0.05 (j: SFC^-^/Veh vs. SFC^+^/Veh).

To exclude the possibility that the extinction-facilitating effects of OXT are mediated by off-target binding to V1aR and V1bR, we next infused a selective OXTR agonist (TGOT) or a specific OXTR antagonist (OXTR-A) bilaterally into LSc. All groups of SFC^+^ mice received the same number of CS-US pairings before they exhibited social avoidance during acquisition of social fear (Fig. 6i). During extinction, all groups showed similar investigation of non-social stimuli, indicating intact exploratory behavior (ns1-ns3; Fig. 6j). However, in SFC^+^/TGOT mice social fear extinction was facilitated, as social investigation levels were significantly higher than that of SFC^+^/Veh mice and similar to that of SFC^-^/Veh mice during the initial phase of extinction (s1-s4; Fig. 6j). In contrast, inhibition of OXTR signaling in SFC^+^/OXTR-A mice impaired extinction as they avoided the presented social stimuli even during the final social exposure (s6; Fig. 6j). Even during recall, social investigation levels of SFC^+^/OXTR-A mice were significantly lower compared to SFC^-^/Veh mice. Finally, TGOT treatment increased, whereas OXTR-A treatment decreased, the proportion of Res mice within their respective SFC^+^ groups compared with SFC^+^/Veh mice (Fig. 6k-o). These results of TGOT-induced facilitation and OXTRA-induced impairment of social fear extinction confirm the importance of OXTR-mediated signaling in the LSc for social valence updating during extinction.

### OXTR-Gαi signaling drives the updating of social valence during extinction

How does enhanced OXTergic signaling within the LSc of Res mice during successful extinction translate to inhibition of LSc^OXTR^ neurons during social contact in Res mice (Fig. 2e, m)? Low levels of OXT activate Gαq signaling downstream of OXTR, while OXT at higher levels activates Gαi signaling ^16^. Interestingly, the physiological effects of Gαi signaling include inhibition of neuronal firing along with reduced neuronal Ca^2+^ influx ^32, 33^. Based on these data and our own findings, we hypothesized that OXT facilitates the reversal of social fear by reducing Ca^2+^ responses in LSc^OXTR^ neurons via Gαi signaling.

To test this hypothesis, we first augmented Gαi signaling downstream of OXTR in the LSc using Atosiban (Ato), which is a biased agonist of OXTR-Gαi signaling (Fig. 7b). Groups of SFC^+^ mice with bilaterally implanted guide cannulas above the left and right LSc were exposed to SFC acquisition (day 1), when they received similar numbers of CS-US pairings to induce social avoidance (Fig. 7c). On day 2, 10 min after pre-extinction intra-LS infusion, all mice exhibited similar investigation of the non-social stimuli (ns1-ns3; Fig. 7d). However, mice that reveived Atosiban (SFC^+^/Ato) mice exhibited facilitated extinction as evidenced by higher social investigation in comparison to the control (SFC^+^/Veh) group during exposure to s2-s4 (Fig. 7d). Even during recall, SFC^+^/Ato mice exhibited enhanced social investigation compared to their SFC^+^/Veh counterparts (s8-s12; Fig. 7d), indicating robust memory formation during extinction. In congruence, the proportion of Res mice was significantly higher in SFC^+^/Ato mice in comparison to SFC^+^/Veh mice (Fig. 7e-g), suggesting that specific activation of OXTR-Gαi signaling promotes association of social stimuli with a positive valence during extinction.

**Fig. 7:**
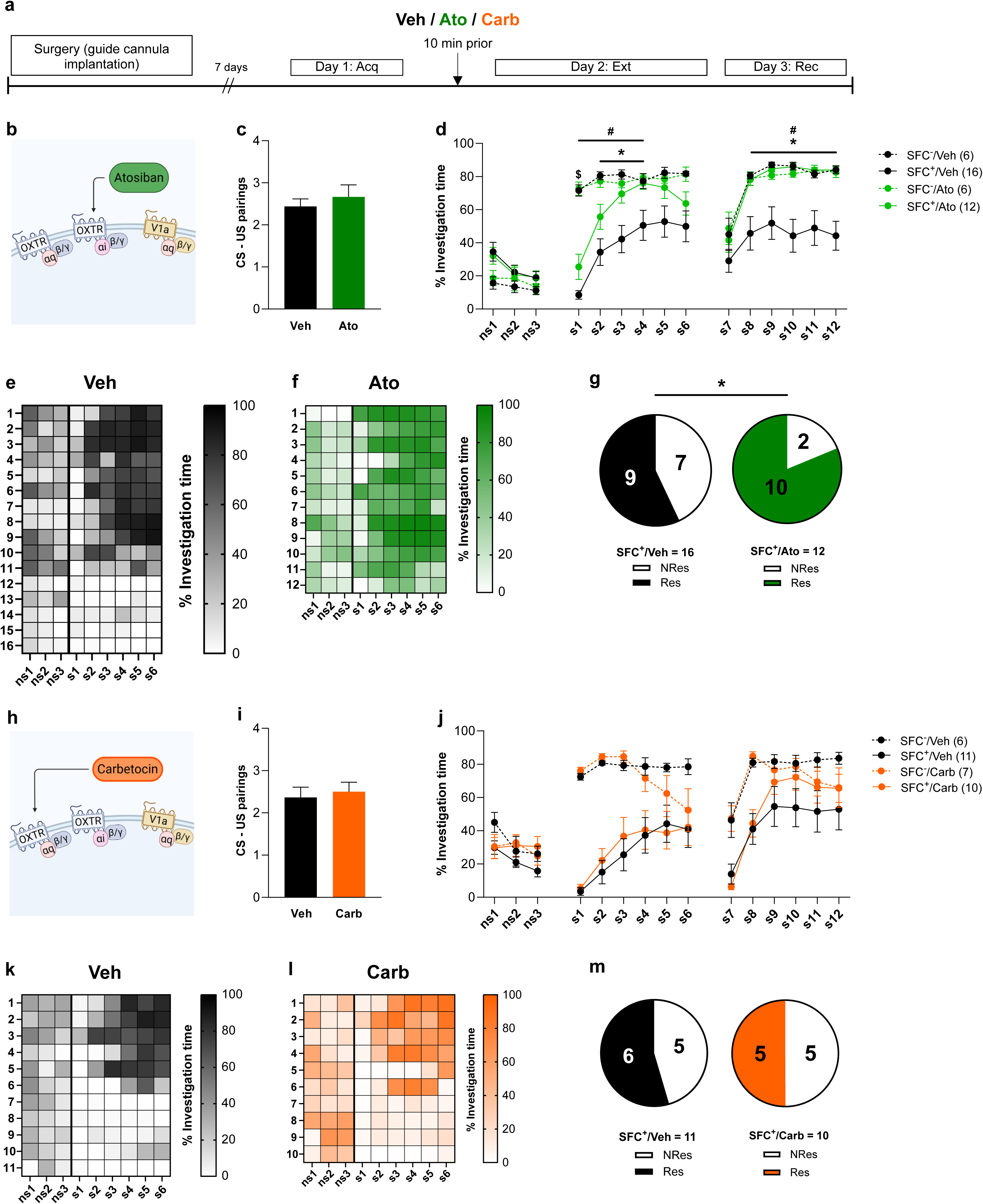
Effect of pharmacological activation of Gαi- or Gαq-coupled signaling downstream of oxytocin receptors (OXTRs) in the caudal lateral septum (LSc) on social fear extinction. **a**, Experimental timeline. **b**, Scheme of Atosiban (Ato; Selective OXTR-Gαi agonist) binding to OXTR. **c**, Number of conditioned – unconditioned stimulus (CS-US) pairings received by social fear-conditioned (SFC^+^) mice during social fear acquisition (Acq). **d**, Social investigation times during social fear extinction (ns1-s6; Ext) and recall (s7-s12; Rec) from SFC^+^ and unconditioned (SFC^-^) mice after bilateral infusions of Ato (5ng/0.2µl/side) or vehicle (Veh; Ringer; 0.2µl/side) 10 min prior to Ext. **e, f**, Heatmap showing the percentage of social investigation of SFC^+^/Veh (e) and SFC^+^/Ato (f) mice during Ext. **g**, Pie chart representing the number of non-responder (NRes) and responder (Res) mice in SFC^+^/Veh vs. SFC^+^/Ato mice. **h**, Scheme of Carbetocin (Carb; Selective OXTR-Gαq agonist) binding to OXTR. **i**, Number of CS-US pairings received SFC^+^ mice during Acq. **j**, Social investigation times during Ext (ns1-s6) and Rec (s7-s12) from SFC^+^ and SFC^-^ mice after bilateral infusions of Carb (5ng/0.2µl/side) or Veh (Ringer; 0.2µl/side) 10 min prior to Ext. **k, l**, Heatmap showing the percentage of social investigation of SFC^+^/Veh (k) and SFC^+^/Carb (l) mice during Ext. **m**, Pie chart representing the number of NRes and Res mice in SFC^+^/Veh vs. SFC^+^/Carb mice. Data represent mean CS-US pairings ± S.E.M (c, i), mean percentage investigation time ± S.E.M (d, j), and absolute number or NRes and Res mice (g, m). *p≤0.05 (d: SFC^+^/Veh vs. SFC^+^/Ato, g), #p≤0.05 (d: SFC^-^/Veh vs. SFC^+^/Veh), $p≤0.05 (d: SFC^-^/Ato vs. SFC^+^/Ato).

We then specifically activated OXTR-Gαq signaling in the LSc using the OXT agonist carbetocin (Carb) (Fig. 7h). However, infusion of carbetocin (SFC^+^/Carb) did not induce difference in the time investigating the non-social stimuli (ns1-ns3; Fig. 7j), the expression of social fear at the start of extinction (s1; Fig. 7j), social investigation times throughout extinction (s1-s6; Fig. 7j), or social investigation times during recall (s7-s12; Fig. 7j) compared to control mice (SFC^+^/Veh). Further, Carb-mediated enhancement of OXTR-Gq signaling did not alter the proportion of Res mice (Fig. 7k-m). This suggests that activating OXTR-Gq signaling in the LSc does not mediate the positive effects of OXT in social valence regulation during extinction.

## Discussion

Successful recovery from social anxiety disorder during exposure therapy requires the reassociation of social cues with positive emotional valence. Using the SFC paradigm, we show that LSc^OXTR^ neurons are indispensable for the aforementioned adaptive social learning. In vivo Ca^2+^ imaging evidenced that LSc^OXTR^ neurons are generally inhibited during social contact, an effect that was lost in SFC^+^ mice, after the acquisition-induced association of negative emotional valence with social contact. Interestingly, restoration of LSc^OXTR^ neuronal inhibition was crucial for successful social fear extinction as seen in Res mice. Integration of multiple imaging features into a composite neuronal score further revealed that the degree of inhibition during social contact, rather than global activity changes, served as the most reliable predictor of extinction success, effectively distinguishing Res from NRes mice. Moreover, we also found that Res mice exhibit higher OXTergic innervation of the LSc relative to the LSr. Pre-extinction chemogenetic silencing of LSc^OXTR^ neurons prevented recovery from social fear, demonstrating the necessity of their specific activity pattern for successful extinction. Finally, pharmacological activation of OXTR-Gi signaling by Ato accelerated extinction and increased the proportion of mice that recovered from social fear, while selective engagement of local OXTR-Gq signaling by Carb had no behavioral effect. Collectively, these findings establish that successful extinction of social fear requires a temporally precise, Gi signaling-mediated inhibition of LSc^OXTR^ neurons during social contact.

Our analyses with receptor autoradiography (Fig. 1b) and OXTR-Cre:*Ai9* reporter mice (Fig. 1c) identified a striking rostro-caudal increase in OXTR expression within the LS. Such sub-regional concentration of OXTRs suggests a functional stratification of the LS, with the LSc mediating the previously identified role for LS-OXT signaling in promoting social fear extinction ^6, 10^. Several recent studies have found that neuroanatomically distinct LS neuronal subpopulations regulate specific adaptive responses to aversive stimuli in social and non-social contexts ^31, 34–39^. Thus, we hypothesized that LSc^OXTR^ neurons may represent a distinct neuronal ensemble involved in the extinction of social fear. Indeed, single-cell Ca^2+^ imaging revealed that LSc^OXTR^ neurons are activated only during investigation of a conspecific in SFC^+^ mice at a time when the social stimulus is associated with a negative valence (s1; Fig. 1j, l). In contrast, these neurons remained largely silent when mice interacted with social stimuli either prior to acquisition (SE; Fig. 1j, k) or after successful fear extinction (s6; Fig. 1j, m). Thus, the excitation of LSc^OXTR^ neurons during s1 suggests that they might promote a negative perception of aversive social stimuli. The main excitatory inputs onto LSc^OXTR^ neurons arise from the ventral hippocampus ^40^. The ventral hippocampus-LS circuit is known for its role in regulating social novelty preference ^41^, driving avoidance in situations presenting an approach-avoidance conflict ^42^ and storing social memory ^43^. Thus, the excitation of LSc^OXTR^ neurons during s1 could result from enhanced hippocampal glutamatergic drive, signaling social fear memory.

Differentially active neuronal subpopulations, including those expressing corticotropin-releasing hormone and dopamine receptors within the LS, regulate responses to aversive social stimuli ^31, 39, 44^. We found that heterogeneous behavioral responses to social fear extinction, as seen in Res and NRes mice (Fig. 2a-e), coincided with distinct activity patterns of LSc^OXTR^ neurons (Fig. 2f, g). Although LSc^OXTR^ neuronal activity patterns were comparable between Res and NRes groups during SE and s1 sessions, only Res mice exhibited the social investigation-associated neuronal inhibition at the end of extinction (s6; Fig. 2f, g). However, social behaviors are dynamic ^45^, and identifying the precise time point of differential activity is essential for dissecting the neuronal mechanisms underlying behavioral variability during extinction. Further sub-second temporal dissection of each social investigation bout revealed that within the SE (Fig 2j, k) and s6 (Fig 2o) sessions, the majority of LSc^OXTR^ neurons were inhibited specifically during the contact phase. This temporal precision strongly suggests that LSc^OXTR^ neurons do not merely encode a global fear state, but rather function as a real-time valence detector that dynamically updates its activity during the most behaviorally relevant epoch of social investigation – the social contact itself. The construction of a composite neuronal score (Fig. 3f) from mean contact activity (Fig. 3c), post-contact increase (Fig. 3d), and contact modulation coefficient (Fig. 3e) confirmed that the degree of contact-associated inhibition was the most robust predictor of extinction success. Notably, the fact that a single integrated score could dichotomize Res and NRes mice highlights that extinction outcome is not determined by a binary switch, but by a continuous spectrum of neuronal response dynamics that collectively shapes the behavioral trajectory during extinction training. Thus, the silent state of LSc^OXTR^ neurons during social contact emerges as a critical neuronal signature of adaptive valence updating, and its absence in NRes mice may represent a failure to engage the inhibitory machinery required to overwrite the traumatic social memory. Thus, inhibition of LSc^OXTR^ neurons during social contact predicted extinction success. Interestingly, we identified a higher OXTergic innervation in the LSc of Res mice (Fig. 4j), a finding that is consistent with our previous data of elevated OXT release in the LS in mice with reduced social fear [6, 10, 11]. Thus, upregulation of OXT signaling in the LSc seems essential for successful social fear extinction, further supporting a prosocial role for OXT ^6, 10, 46–48^.

However, this raises an important question: how does enhanced OXT release in the LSc lead to reduced social contact-associated activity of LSc^OXTR^ neurons, as seen during s6 and in Res mice? Two distinct neuronal mechanisms could explain such counterintuitive findings. Firstly, feedforward inhibition, which is a feature of the LS with the majority of neurons being GABAergic, could underlie the observed inhibition of the LSc^OXTR^ neurons ^9, 49^. Secondly, the enhanced involvement of Gαi-GPCR signaling downstream of OXTR could result in inhibition of LSc^OXTR^ neurons upon OXT stimulation. In vitro studies have shown that higher concentrations of OXT activate Gαi-coupled GPCR signaling downstream of OXTR ^16, 50^. Hence, an upregulation of OXT signaling, resulting from increased OXT-positive fiber density in Res mice (Fig. 4j) and enhanced LS-OXT release in mice expressing reduced social fear, could bias OXTR coupling to Gαi proteins, thereby inhibiting LSc^OXTR^ neurons during positive social contact.

Specific chemogenetic silencing of LSc^OXTR^ neurons prior to extinction impaired extinction and increased the number of NRes mice within the SFC^+^/CNO group (Fig. 5d-g). This result confirms the necessity of LSc^OXTR^ neurons for successful social fear extinction. Taken together with our previous data, it also suggests that it is not merely the presence or absence of LSc^OXTR^ neuronal activity that matters, but rather the precise temporal structure of their activity patterns during the contact phase. The composite neuronal score derived from our Ca²⁺ imaging data demonstrated that successful extinction in Res mice is phenocoded by transient inhibition of LSc^OXTR^ neurons specifically during social contact. Disruption of this pattern by global chemogenetic silencing of LS^OXTR^ neurons impaired extinction and recapitulated the phenotype observed in NRes mice. This suggests that LSc^OXTR^ neurons are not merely permissive for extinction, but actively orchestrate valence reassignment through their temporally defined inhibition during social contact.

Causal evidence for the specific involvement of Gαi-GPCR signaling downstream of OXTR comes from our pharmacological experiments. Activating OXTRs in the LS with infusions of synthetic OXT facilitated the extinction of social fear (Fig. 6c-h). This effect of OXT is mediated via OXTR signaling, as TGOT infusion facilitated social fear extinction, whereas OXTR-A infusion impaired it (Fig. 6l-q). These data rule out the possibility that the behavioral effects of OXT are due to cross-binding to V1aR and V1bR, which are abundantly expressed in the LS ^51^. Additionally, these data corroborate the results of our chemogenetic experiment, suggesting that endogenous OXT signaling is essential for successful extinction. Both synthetic OXT and TGOT lead to complete OXTR activation, i.e., they activate both Gαq and Gαi signaling downstream of OXTR ^16, 52^. Following this, agonizing LSc^OXTR^ with Ato (OXTR-specific Gαi agonist) dramatically reduced social fear expression, facilitating extinction (Fig. 7d-g), whereas Carb (OXTR-specific Gαq agonist) had no effect in this context (Fig. 7j-m). On a circuit level, Ato-mediated activation of Gαi signaling still allows LSc^OXTR^ neurons to participate in circuit dynamics driven by non-OXT inputs. In contrast, chemogenetic inhibition of LSc^OXTR^ neurons completely abolishes their influence on septal circuitry. Taken together with our earlier findings, our data propose that activation of OXTR-Gαi by natural variation in the OXTergic innervation of LSc in Res mice drives their extinction success.

The findings of this study can be extrapolated to a broader conceptual framework linking neuronal temporal dynamics to the adaptive regulation of social valence. Naturally higher OXT availability in the LSc, due to increased release during positive social contact, promotes the silencing of LSc^OXTR^ neurons via OXTR-Gαi signaling. Thus, OXTR-Gαi signaling acts as a neuronal brake, inhibiting social fear memory and promoting the social valence reassignment, thereby allowing for fine-tuned adaptation of responses to aversive social cues. These findings establish a mechanistic framework linking individual variation in OXT-dependent regulation of LSc^OXTR^ neuronal dynamics to extinction success, with potential relevance for variability in response to exposure-based therapy in social anxiety disorder.

## Acknowledgements

This work was supported by the German Research Foundation (DFG) grant (ME5731/2 to RM; Ne465/31 to IDN; CRC1089 to TK), the DFG Graduate School (GRK-2174; to IDN and RM), and the Alexander von Humboldt Foundation and DFG Walter Benjamin Program (project number 551841466; to DK).

## Author contribution statements

Project conception: RM and IDN; Methodology: TS, RM, FS, DK, AB, TK, and IDN; Anatomic analysis: LB, RM; In vivo calcium imaging: TS, FS, and DK; Behavior analysis: TS, AK, and RM; Writing: RM, TS, DK, TK, and IDN; Illustrations: RM, TS, and DK; Project administration and supervision: TK, IDN, and RM.

## Supplementary Figures

**Fig. S1:**
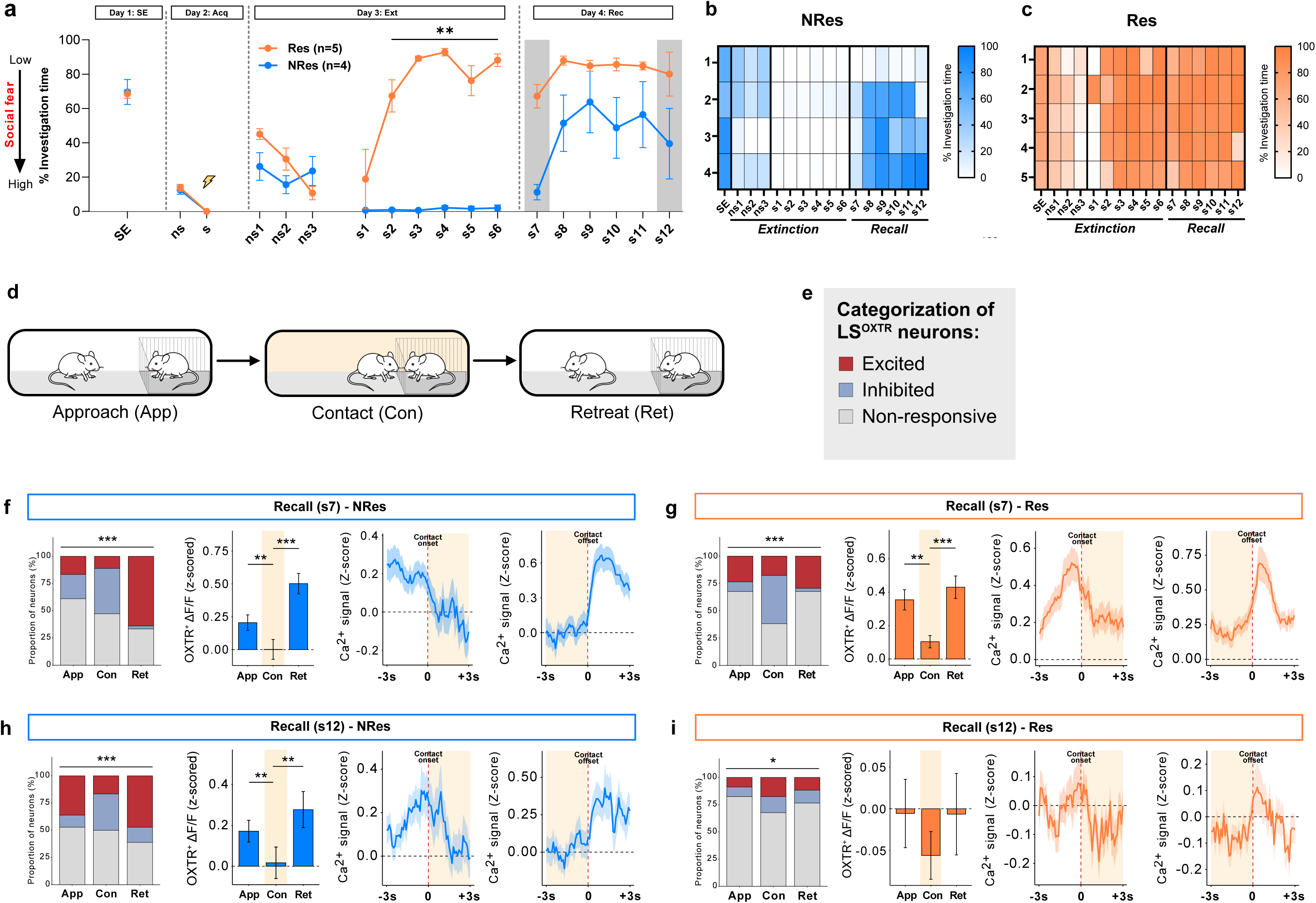
Activity dynamics of oxytocin receptor (OXTR) expressing neurons in the caudal lateral septum (LSc^OXTR^) during the recall phase of the social fear conditioning (SFC) paradigm in non-responder (NRes; Blue) and responder (Res; Orange) mice. **a**, Percentage of time mice spent investigating non-social (ns) and social (s) stimuli in the SFC paradigm (SE: Social exposure; Acq: Acquisition; Ext: Extinction; Rec: Recall) in NRes and Res mice. **b, c,** Heatmaps showing the percentage of time spent with social investigation of NRes (b) and Res (c) mice during SE, Ext, and Rec. **d,** Schematic representation of the temporal stratification of a single social investigation bout into approach (App), contact (Con), and retreat (Ret) phases. **e,** Legend for the categorization of LSc^OXTR^ neurons. **f-i,** Percentage of excited (red), inhibited (light blue), and non-responsive (grey) LSc^OXTR^ neurons of NRes (f, h; n = 59 and 68 neurons, respectively) and Res (g, i; n = 49 and 49 neurons, respectively) mice along with a representative neuronal trace during s7 (Day 4; f, g) and s12 (Day 4; h, i) sessions. Data represent mean percentage investigation ± S.E.M (a) and percentage of neurons (panel 1), mean OXTR ΔF/F ± S.E.M (panel 2), and Ca^2+^ signal Z-score (panel 3 and 4) (j-o). *p≤0.05 (i), **p≤0.01 (a, f, g, h), ***p≤0.001 (f, g, h).

**Fig. S2:**
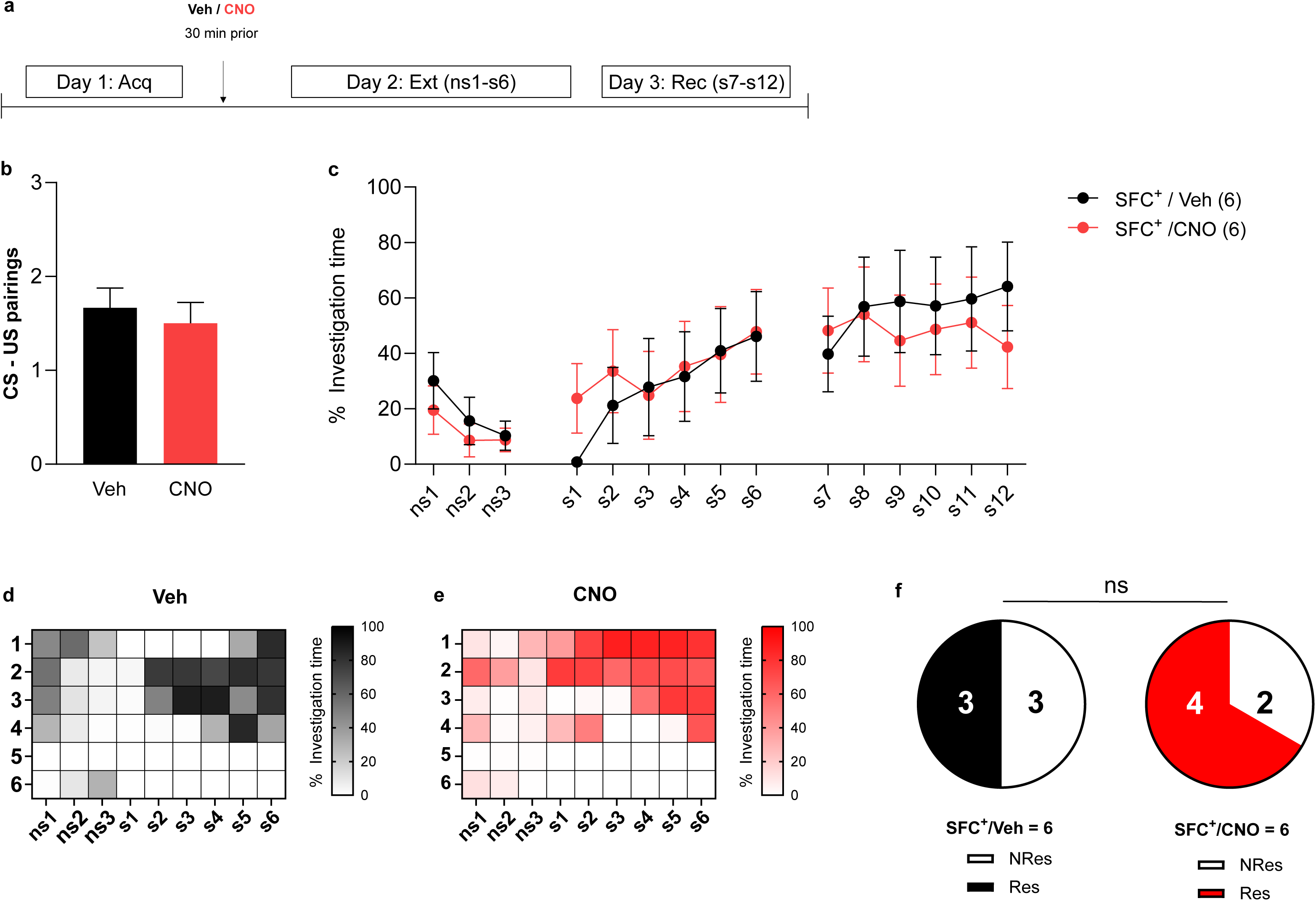
Effect of CNO on social fear extinction. **a**, Timeline for the experiment including pharmacological manipulation and social fear conditioning (SFC) paradigm (Day 1-3). **b**, Number of conditioned-unconditioned (CS-US) stimulus pairings received by social fear conditioned (SFC^+^) mice during social fear acquisition (Acq). **c**, Social investigation times during social fear extinction (ns1-s6; Ext) and recall (s7-s12; Rec) of SFC^+^ and unconditioned (SFC^-^) mice treated with either clozapine-n-oxide (CNO; 10 mg/kg/0.2ml of saline; i.p.) or vehicle (Veh; 0.2ml of saline; i.p.) 30 min prior to extinction. **d, e,** Heatmaps showing the percentage of social investigation of SFC^+^/Veh (e) and SFC^+^/CNO (f) mice during Ext. **f**, Pie chart representing the number of non-responder (NRes) and responder (Res) mice within SFC^+^/Veh and SFC^+^/CNO groups. Data represent mean CS-US pairings ± S.E.M (b), mean percentage investigation time ± S.E.M (c), and absolute number of NRes and Res mice (f).

## Notes

### Competing Interest Statement

The authors have declared no competing interest.

